## Supplemental Figures for "HDX-MS reveals structural determinants for RORγ hyperactivation by synthetic agonists"

**Supplemental materials**

**SUPPLEMENTAL FIGURE LEGENDS**

**Supplementary Figure 1: Overview of HDX-MS analysis of RORγ**. **A**) Sequence coverage map of RORγ resulting from in-line pepsin digest. Representative deuterium incorporation curves for helix 12, helix 3, helix 7, and helix 3, are shown in panels **C**, **D**, **F**, and **G**, respectively. Results for differential HDX-MS expereiments showing the effects of coactivator peptides are shown in panels **B** and **E**. RORγLBD + SR19265 vs RORγLBD +SR19265 + SRC1-2 is shown in **B**. RORγLBD + SR19265 + SRC1-2 vs RORγLBD-SRC2 fusion construct are shown in panel **E**.

**Supplementary Figure 2: Ligand electron density for reported co-crystal structures.** Composite omit map densities were generated with simulated annealing and the density corresponding to the ligand are overlaid with the ligand model. SR19265 and SR19355 are shown in panels **A** and **B**, respectively. SR2211 was found in both molecules in the asymmetric unit and the densities for the ligands of chain A and chain C are shown in panels **C** and **D**, respectively.

**Supplementary Figure 3: Reproducibility and correlation of Biochemical, biophysical and HDX-MS measurements**. AlphaScreen peptide recruitment and differential scanning fluorimetry based thermal stability assays were tested on separate days with two batches of protein and the results are compared in figures **A** and **B**. Each data point represents a single compound and SR19355 was found to be an outlier for the thermal stability assay. The distribution of compound performance for both assays are shown in panels **C** and **D**. AlphaScreen activity was compared to affinity (EC50) for fluorescent peptides as determined by fluorescence polarization. The results from a correlation analysis are shown in panel **E**. The reproducibility of differential HDX-MS two-timepoint screening was assessed by comparing perturbation values (Δ%D) of a subset of 16 modulators (data unpublished) in panel **F**. The results from a weighted Pearson correlation analysis comparing function assay data from AlphaScreen-based peptide recruitment, ligand dependent thermal stability, and cell-based activity are shown in panel **G**. P values that were lower than 0.05 were colored in red and R^2^ values greater than 0.4 were colored in green.

**Supplementary Figure 4: Results of covariation analysis**. A weighted correlation analysis of the two timepoint compound screening dataset shows widespread correlation between peptides. Across the diagonal are histograms showing normal distributions of Δ%D for each peptide. Below the diagonal are correlation plots comparing peptide perturbation values of the peptides above and on the right of the corresponding plot. Above the diagonal are correlation coefficients (Pearson’s R value) of a linear model describing the variances of peptide perturbation values of the peptides below and left of the corresponding R value. Significance is indicated for adjusted P vales < 0.1, 0.05, 0.01 and 0.001 as ., *,**, and *** respectively.

**SUPPLEMENTAL FIGURES**

**Figure 1**

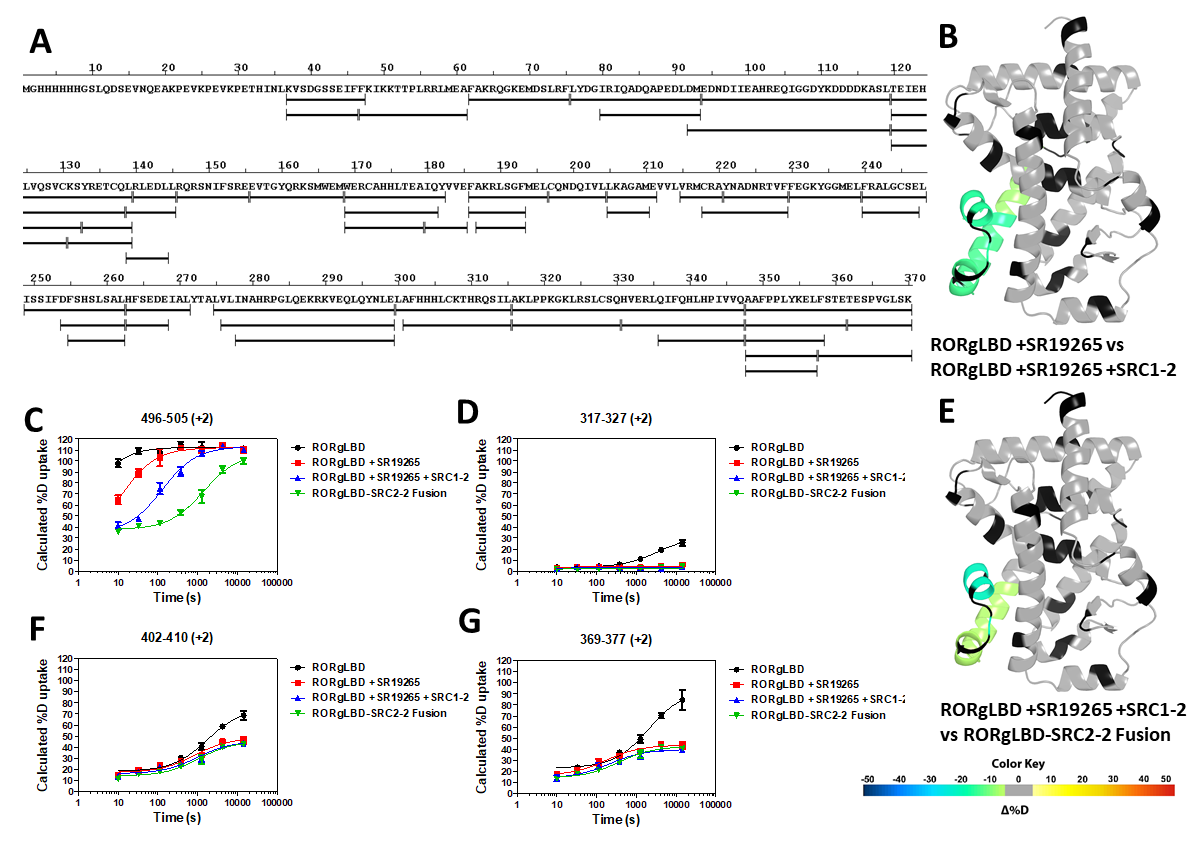

**Figure 2**

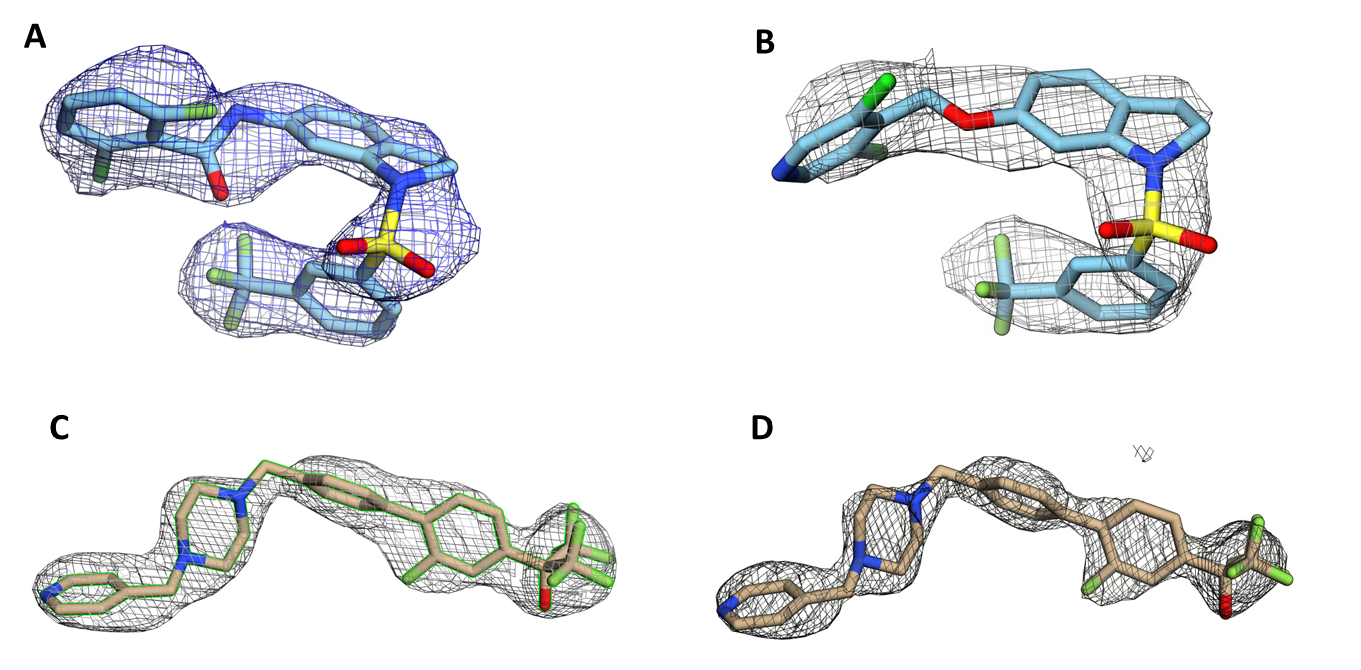

**Figure 3**

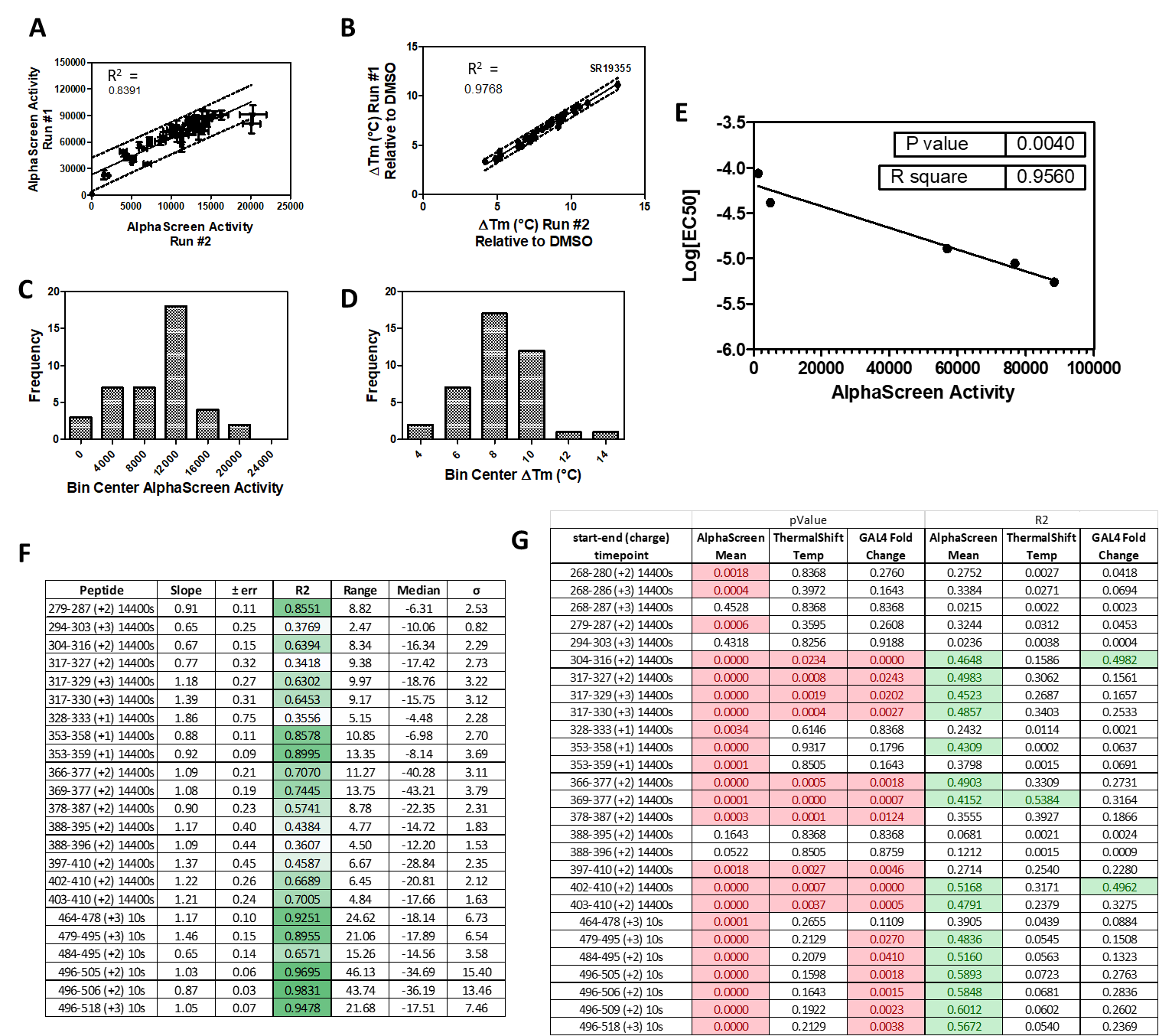

**Figure 4**

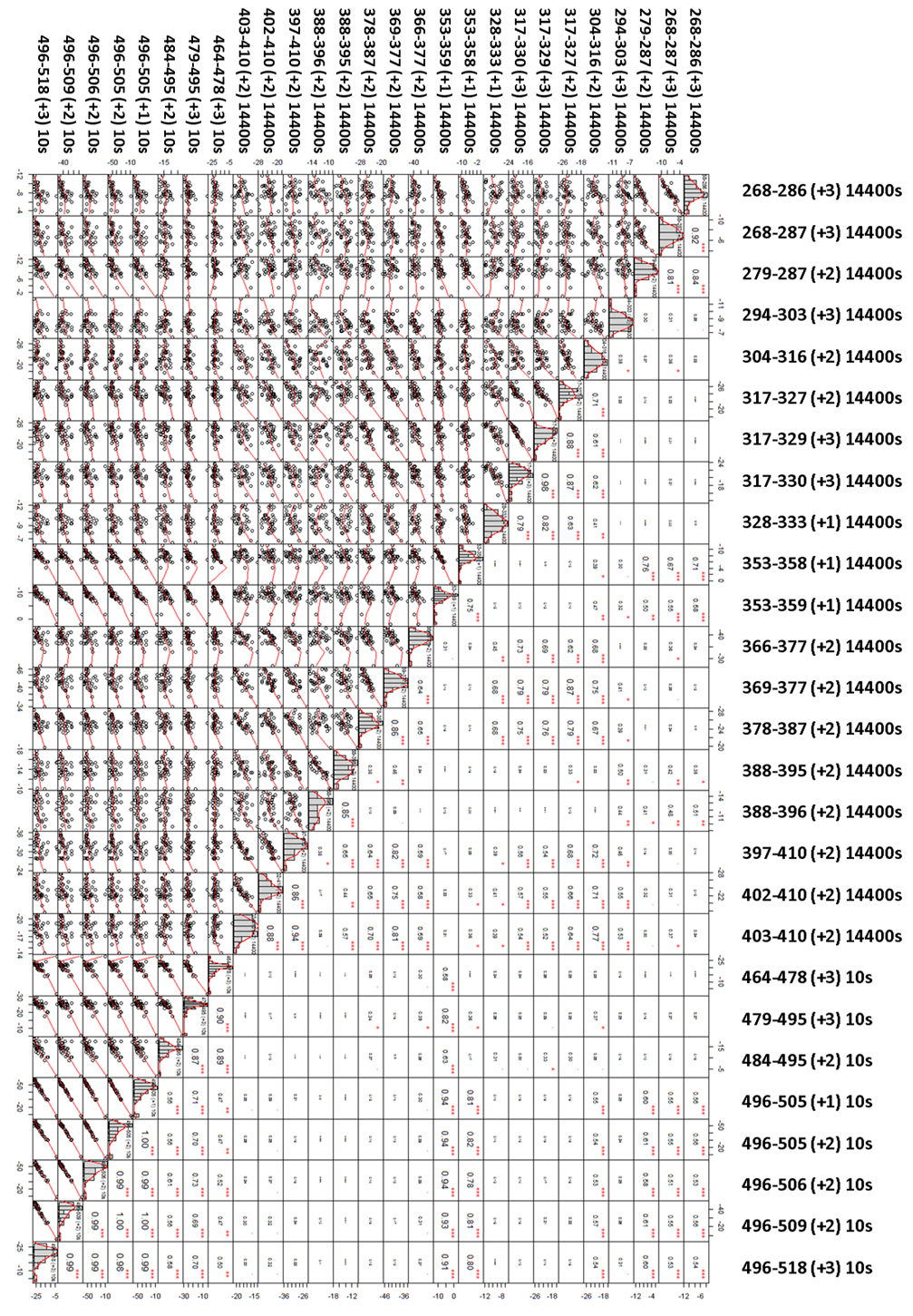
